# Cochlear nucleus small cells use olivocochlear collaterals to encode sounds in noise

**DOI:** 10.1101/2021.05.20.444983

**Authors:** Adam Hockley, Calvin Wu, Susan E Shore

## Abstract

Understanding communication signals, especially in noisy environments, is crucial to social interactions. Yet, as we age, acoustic signals can be disrupted by cochlear damage and the subsequent auditory nerve fiber degeneration. The most vulnerable—medium and high-threshold—auditory nerve fibers innervate various cell types in the cochlear nucleus, among which, the small cells are unique in receiving this input exclusively. Furthermore, small cells project to medial olivocochlear (MOC) neurons, which in turn send branched collaterals back into the SCC. Here, we use single-unit recordings to characterise small cell firing characteristics and demonstrate superior intensity coding in this cell class. We show converse effects when activating/blocking the MOC system, demonstrating that small-cell unique coding properties are facilitated by direct cholinergic input from the MOC system. Small cells also maintain tone-level coding in the presence of background noise. Finally, small cells precisely code low-frequency modulation more accurately than other ventral CN (VCN) cell types, demonstrating accurate envelope coding that may be important for vocalisation processing. These results highlight the small cell–olivocochlear circuit as a key player in signal processing in noisy environments, which may be selectively degraded in aging or after noise insult.

## INTRODUCTION

Auditory perception in ever-changing soundscapes is key for communication. However, this important task is degraded with age (Pichora-Fuller, 1997). Crucial to acoustic signal processing are the temporal fine structure and envelope information present in complex signals (Shannon et al., 1995), and difficulties in resolving these signals by the brain are prevalent in age-related hearing deficits (Bharadwaj et al., 2014). In the auditory periphery, the high threshold, low spontaneous rate (SR) auditory nerve fibers (ANFs) may be particularly important as their best dynamic range overlaps with normal levels of communicative signals. These specialised ANFs project along with low threshold ANFs throughout the cochlear nucleus (CN). However, the small-cell cap region receives exclusive input from the low (and medium) SR ANFs (Liberman, 1991; Ryugo, 2008).

Located between the granule-cell and magnocellular domains of the VCN (Cant, 1993), the small-cell cap (SCC), as the name suggests, is a region containing small cells (<20 μm). In humans, the SCC occupies a larger proportion of the CN than in rodents, and is therefore poised to play a major role in central mechanisms of speech perception (Moore & Osen, 1979). Small cells project either to medial olivocochlear (MOC) neurons in the ventral nucleus of the trapezoid body (VNTB; Darrow et al., 2012; De Venecia et al., 2005; Thompson & Thompson, 1991; Ye et al., 2000) or directly to the medial geniculate body, the latter serving as short-latency relay to the auditory cortex (Schofield, Mellott, et al., 2014; Schofield, Motts, et al., 2014). Small cells, in turn, receive branched collaterals from MOC neurons (Benson et al., 1996; Benson & Brown, 1990; Ryan et al., 1990), whose major function is suppression of cochlear outer hair cell electromotility to alter cochlear gain. MOC activation releases the cochlea from background masking by shifting its dynamic range (Dolan & Nuttall, 1988; Kawase & Liberman, 1993). However, small cells remain largely unexplored. Here, we hypothesise that MOC collateral projections to the VCN, and especially onto small cells, may counteract the effect of cochlear suppression, allowing small cell firing to accurately represent stimulus intensity.

In this study, we characterised small cell response properties that are consistent with its unique circuit arrangement. We showed that small cells can more accurately encode intensity as well as temporal information compared to other VCN cell types. Further, we found both MOC activation and inhibition of cholinergic input to the VCN bidirectionally modulated small cell firing properties. In the presence of background noise, small cells maintained superior intensity coding. Taken together, these findings reveal a special role for small cells in processing communicative signals and that its degradation after selective auditory neuropathy may underlie hearing-in-noise difficulties.

## METHODS

All animal experimental procedures were performed in accordance with the protocols established by the National Institutes of Health (Publication 80-23) and approved by the University Committee on Use and Care of Animals at the University of Michigan. Male and female (n = 11 and 15, respectively) pigmented guinea pigs weighing 280-800g obtained from Elm Hill Labs were used. No differences in unit responses for animal’s sex or age was found. Guinea pigs were dual-housed on a 12/12 h light-dark cycle, with food and water readily available. Auditory brainstem responses (ABRs) were used to confirm normal auditory thresholds of ≤20 dB SPL at 4-20kHz in all animals. ABRs (0–90 dB SPL tone bursts; 5 ms duration, 1 ms rise/fall times, 21 Hz presentation rate, 512 repetitions in 10 dB steps; Tucker-Davis Technologies RZ6) were recorded at 4, 8, 12, 16 and 20 kHz.

### Surgery

Guinea pigs were initially anaesthetised with ketamine (50 mg kg^−1^; Hospira Inc., Lake Forrest, IL, USA) and xylazine (5 mg kg^−1^; Akorn Inc.; Lake Forrest, IL, USA). Anaesthetic depth was maintained using 10 mg kg^−1^ ketamine and 1 mg kg^−1^ xylazine supplements. Atropine (0.05 mg kg^−1^) was administered during the initial surgery to reduce bronchial secretions. Animals were placed in a stereotaxic frame (Kopf; Tuijunga, U.S.A.) within a sound-attenuating and electrically shielded double-walled chamber. The skull was exposed, and a midline incision made, temporalis muscle removed, and a craniotomy performed to allow access to the left CN. The dura mater was removed and the exposed brain surface was kept moist by regular applications of saline.

### Sound presentation

Auditory stimuli were delivered monaurally via a closed-field, calibrated system (modified DT770 drivers; Beyerdynamic Heilbronn, Germany) coupled to hollow ear bars. The speakers were driven by a Tucker-Davis Technologies (TDT; Alachua, FL, USA) System 3 (RZ6, PA5 & HB7), controlled by TDT Synapse and custom MATLAB 2021a (Mathworks; Natick, U.S.A) software.

### Neural recordings

Multi-channel recording probes (NeuroNexus; Ann Arbor, MI, USA) were advanced stereotaxically through the cerebellum towards the left CN using an MP-285 microdrive (Sutter Instruments; Novato, CA, USA). Signals were amplified by a TDT PZ5 preamp connected to a TDT RZ2 processor for filtering (0.3 – 5 kHz), and data was collected using Synapse software. Spikes were detected on-line with threshold set at 4 standard deviations from mean background noise, then spike-sorted post-hoc using a PCA approach. Electrodes were positioned so that shanks passed through the dorsal cochlear nucleus (DCN) to reach the SCC and VCN. In 2 animals, electrode tracts were marked to confirm SCC positioning. Units were characterised by responses to tone-burst and broadband noise (BBN) signals, also used to create receptive fields, peri-stimulus time histograms (PSTHs) and rate-level functions (RLFs) (Ghoshal & Kim, 1997; Palmer, 1987; Stabler et al., 1996; Winter & Palmer, 1995), and manually characterised divided into 5 categories; PL (bushy cells), Ch (T-stellates), SC (small cells), B (buildups) and On (onsets). A machine learning model was then used to confirm manual typing had produced discrete classifications of neurons (Fig 1). Poorly driven units were discarded.

**Fig 1:**
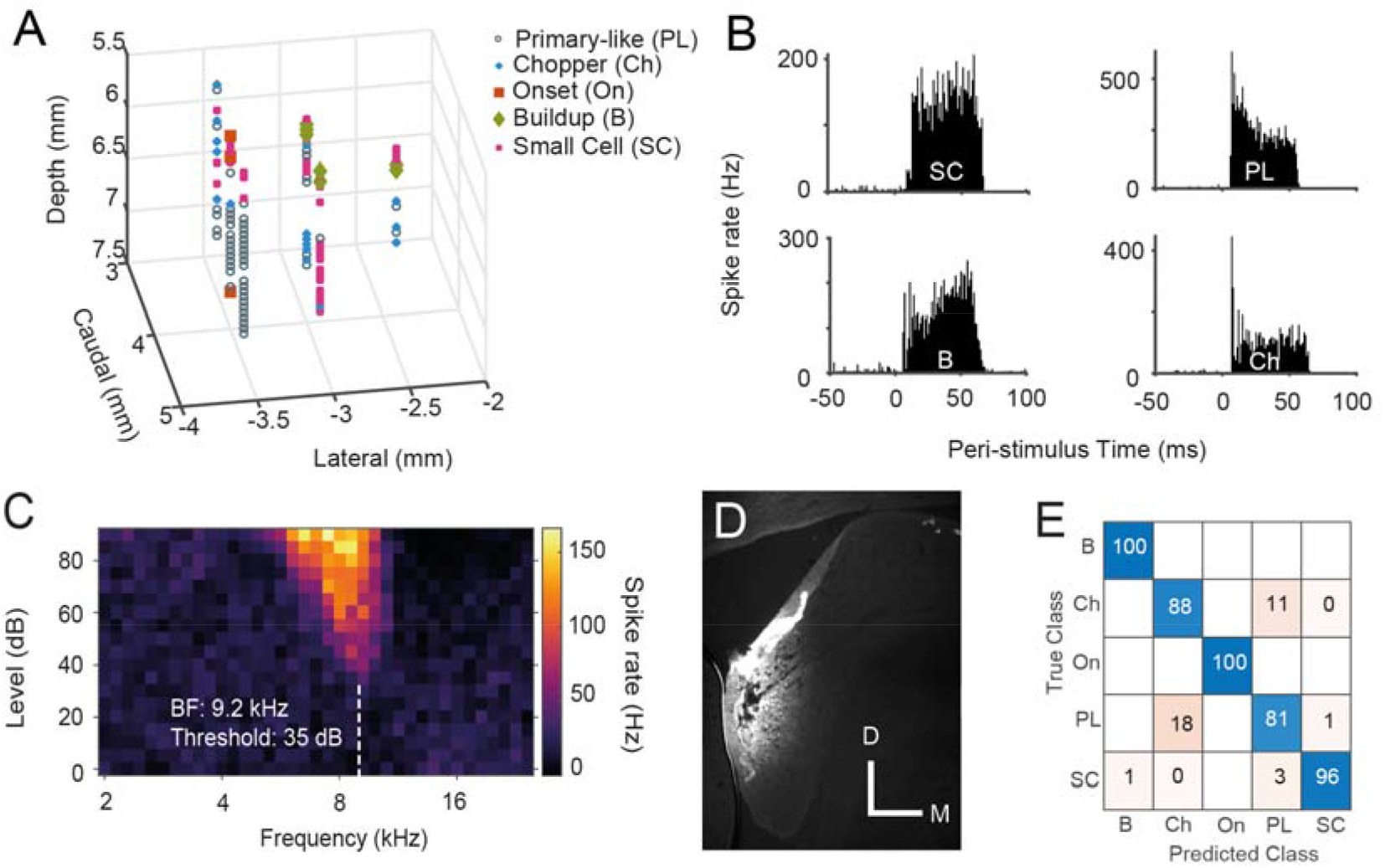
Validating recordings from CN small cells. A) An example 3D plot of unit types within the left CN of one guinea pig. Cells exhibiting buildup PSTH shapes were found more medial and caudal, in the DCN. The VCN is more rostral and lateral, with the SCC units located between these two structures. B) Example PSTHs from 4 different CN cell types: small cell (SC) with flat, bimodal PSTH, primary-like (PL), buildup (B) and chopper (Ch). C) Receptive field of a CN small cell. D) Coronal section of the cochlear nucleus, containing a Fluorogold-marked electrode track in the SCC. (Bar = 0.5 mm) E) Confusion matrix of the machine learning model used to confirm recordings from discrete types of neurons.

### Intensity coding analyses

Receptive field stimuli consisted of randomised tonebursts (50 ms duration, 200 ms ISI, 0.1 octave spacing, 5 dB steps, 15 repeats, randomised). From these, threshold and BF were obtained from the lowest sound level to produce firing >2 standard deviations above spontaneous firing rates. RLFs at BF were then used for further analyses. Total inhibitory area of the receptive field was calculated as the number of squares of the receptive field where firing rate was <2 standard deviations below spontaneous firing rates.

For further intensity coding analysis, a PSTH-based stimulus classification was performed using methods described in Foffani & Moxon (2004). Briefly, spike responses (serialised PSTH; 1 ms bins concatenated across units) of the neural population generated in each trial (50 trials total) were compared to the “template,” or average PSTH, of a given stimulus intensity. Euclidean distances between each trial and the templates (19 total: 0 to 90 dB with 5 dB steps) were computed and the template with the shortest distance was then assigned as “classified intensity.” Classification accuracy was defined as the percentage of trials that were classified within <10 dB of the actual intensity, or “hits.” Coefficient of variation for classification result across all 20–50 trials was calculated as std(x^2^)/mean(x^2^), where x counts the discretised step away from the “hit.”

### Drug delivery

Local drug delivery of the muscarinic acetylcholine receptor antagonist, atropine was achieved using a NeuroNexus D16 puffer-probe. The probe was connected to a 100 μl syringe containing an 80 μM solution. The syringe was fixed to a digital syringe driver (UMP3 microsyringe injector with Micro4 controller; World Precision Instruments; Sarasota, FL). This system allows simultaneous drug delivery and neural recordings as described and validated in previous studies (Rohatgi et al., 2009; Stefanescu & Shore, 2015, 2017). After assessing baseline CN neuronal activity, 2 μl of solution was delivered at a rate of 100 nl/min, as in previous studies (Rohatgi et al., 2009; Stefanescu & Shore, 2015, 2017).

### Round window recordings

Cochlear compound action potential (CAP) recordings were performed using an insulated silver-wire ball inserted through the bulla to rest on the cochlear round window. The trailing wire was sealed to the skull, with a small hole left open for fluid wicking. CAPs were recorded in response to 10 ms tones with rise/fall times of 2 ms. Amplitudes were calculated as P_1_-N_1_ of the mean CAP waveform (200 repeats).

### Electrical stimulation

MOC neurons in the VNTB were electrically stimulated with biphasic current pulses (100 ms duration, 0.1 ms pulse width, 100 Hz, 500-1000 μA; 5/sec) applied through concentric bipolar electrodes (CBCEF75; FHC; Bowdoin, ME). CAPs were recorded to verify MOC neuron location (16 kHz, 70 dB SPL tone). CAP amplitudes were compared at baseline and 5 ms after the cessation of electrical stimulation. A CAP-amplitude reduction of >30% confirmed accurate MOC location in the VNTB for stimulation. Current levels were titrated to ensure maximal MOC activation with no elicited muscular contractions.

### Imaging

In a separate guinea pig, a tract tracing experiment was performed to confirm VNTB projections to and from SCC. FluroEmerald (10%; 0.5 μl, Thermo Fisher Scientific, Waltham, MA, USA) was pressure-injected (0.1 μl min-1) into the contralateral VNTB (1 mm anterior to the interaural line, 1.5 mm lateral to midline, 12 mm from the dural surface). The animal was anesthetised with ketamine-xylazine mixture in the same manner as described for electrophysiology experiments but allowed to recover and maintained in the vivarium for 5 days before transcardial perfusion (100 mL 1× PBS, 400 mL 4% paraformaldehyde) after euthanasia. The brain was extracted, post-fixed for 2 h, immersed in 30% sucrose solution for 2 days, frozen and cryosectioned at 50 μm in the coronal plane (Leica, CM3050S). Coverslipped brain sections were examined under confocal epifluorescence (PMT; Leica, SP5-x).

### Experimental design and statistical analysis

Small cell sound evoked responses were compared to other VCN cell types. Changes in these responses during MOC modulation were also analysed. Statistical tests, including Kruskal-Wallis test, ANCOVA, and two-way ANOVA were used to compare between cell-types or conditions (α = 0.05). Post hoc analyses were performed using Tukey’s multiple comparison test.

## RESULTS

Using multichannel electrodes, we isolated 1523 single units from 83 VCN electrode locations in 26 guinea pigs. Unit distributions are shown in Table 1. We excluded onset and buildup units in subsequent analyses due to their low numbers.

**Table 1:** Number of single CN units recorded. Cell types are Primary-like (PL), Chopper (Ch), Small cell (SC), Onset (On) and Buildup (B).

|  | PL | Ch | SC | On | B | Total |
| --- | --- | --- | --- | --- | --- | --- |
| **Control** | 722 | 247 | 310 | 57 | 187 | 1523 |
| **VNTB stimulation** | 260 | 96 | 139 | 15 | 75 | 585 |
| **Atropine infusion** | 20 | 36 | 34 | 4 | 8 | 102 |

Single-shank 32-channel electrodes with 50 µm spacing of electrode sites were positioned to simultaneously record units from the SCC and VCN (Fig 1A). Units were classified by their temporal and frequency response patterns (Fig 1B)(Stabler et al., 1996; Winter & Palmer, 1995; Young et al., 1988). Small cells, located dorsoventrally between DCN and VCN, displayed several unusual PSTH shapes, the most common being a ‘flat’ or bimodal PSTH shape (Fig 1B). Small cells were therefore identified by a combination of their PSTH shapes and unique locations within the cochlear nucleus and by their distinct best frequency (BF) progressions with depth. A machine learning model (RUSBoosted Trees) was used to confirm that manual typing had produced discrete types of neurons (Fig 1E). This model used 23 inputs, each of which was a metric obtained from either the PSTH, inter-spike interval histogram (ISIH), receptive field or RLF.

To confirm reciprocal projections from small cells to MOC neurons in the VNTB, a tracer (FluoroEmerald) was pressure-injected into the VNTB. One week later, retrograde and anterograde staining was observed in the SCC (Fig 2), demonstrating, along with the physiological data, that SCC neurons form a circuit with MOC neurons, confirming previous findings (Benson et al., 1996; Benson & Brown, 1990; Ye et al., 2000).

**Fig. 2:**
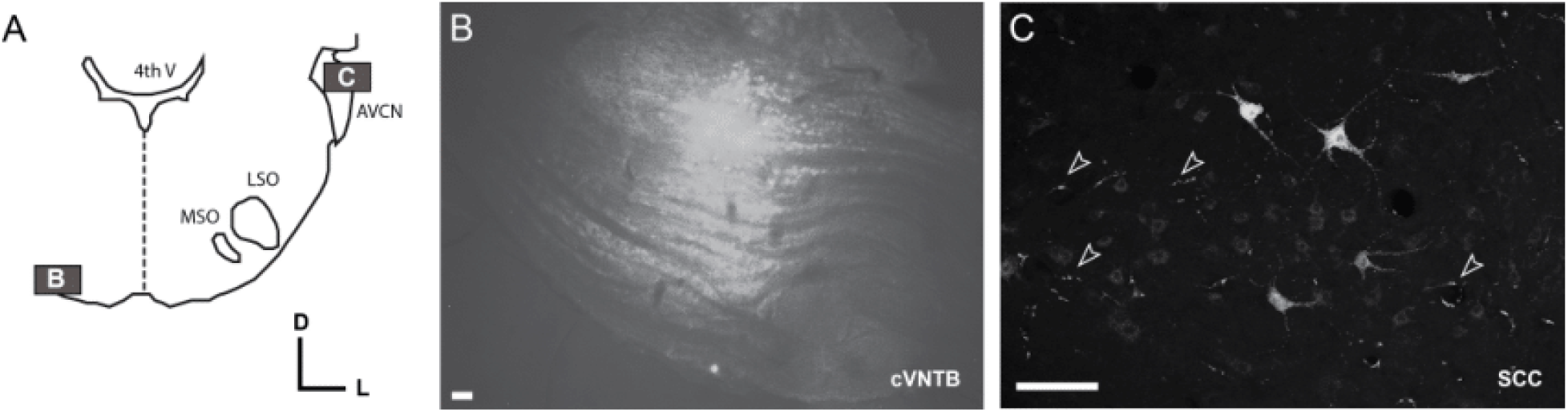
VNTB neurons project to and receive projections from the SCC. A) Schematic showing the locations of photomicrographs in B (VNTB) and C (SCC). B) FluoroEmerald injection in the contralateral VNTB (cVNTB). C) Anterograde labeling of puncta (arrowsheads) in SCC originating in VNTB and retrograde labelling of SCC somata. Bar = 100 µm.

### Small cell firing patterns resemble those of low/medium SR ANFs

Afferent input to small cells is exclusively from low/med SR, high threshold ANFs, while small cells project to, and receive input from MOC neurons. Due to this unique circuit configuration, we hypothesised that small cells would have different patterns of intensity coding (rate-level functions) compared to VCN cell types that also receive input from high SR, low threshold ANFs. small cell response characteristics were like their low/medium SR, high threshold ANF inputs, with low SRs, high thresholds, and long first-spike latencies compared to other VCN cell types (Fig S1). Small-cell receptive fields contained inhibitory areas (Figs 1C S1D), signifying a currently unknown source of inhibition.

To examine intensity coding at speech-relevant levels, we used a PSTH-based classifier to compare the trial to trial reliability across cell types in response to different intensity stimuli (Foffani & Moxon, 2004). Intensity coding fidelity is represented by accurate representations of stimuli for each single-trial spiking response (Fig 3C). Primary-like units displayed better intensity discrimination between 20-45 dB as well as very low variability, while small cells were better at intensity coding above 60 dB (Fig 3D–E; ANCOVA F=9.1, P=1.1e-4, effect size (partial eta squared) = 0.006). The high-fidelity intensity coding and absence of RLF plateaus in small cells is likely due to their combined input from low SR ANFs and excitatory input from MOCs at high sound levels. The dual action of peripheral and central excitation would allow small cells to better encode tone intensities at levels relevant for speech processing.

**Fig 3:**
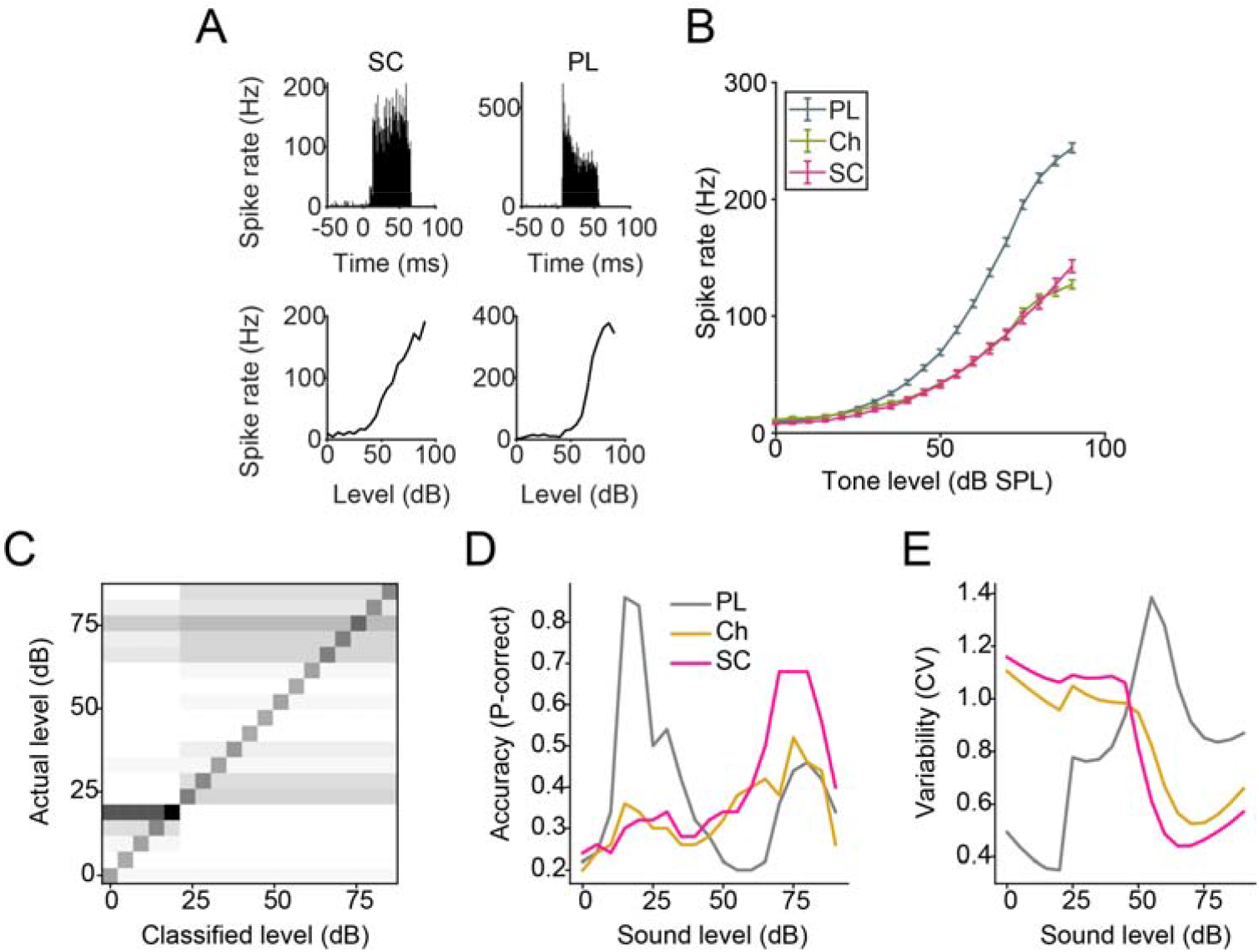
CN small cells show superior intensity coding at high sound levels. A) Example PSTH and RLFs for a CN small cell and a primary-like unit. B) Mean (± SEM) RLFs of 4 major CN cell types. C) Confusion matrix for sound intensity classification across the population (all cell types). Greyscale – white pixels: 0%, black pixels: 100%. D) Probability of correct classification across unit-types. E) Trial to trial variability (coefficient of variation) of classifier performance across unit-types.

### MOC neurons modulate small-cell excitability

Cholinergic MOC neurons send projections into the SCC (Benson et al., 1996; Benson & Brown, 1990; Ryan et al., 1990). Acetylcholine is known to act on CN cells through muscarinic receptors, resulting in a cell-type specific excitation or modulation of neural plasticity (Fujino & Oertel, 2001; Stefanescu & Shore, 2017). To investigate whether cholinergic MOC inputs mediate the greater accuracy of intensity coding in small cells, first, we applied the muscarinic acetylcholine receptor (mAChR) antagonist, atropine using drug delivery probes. Atropine reduced small cell medium to high intensity tone-evoked activity, (e.g. for 75 dB SPL, α = 0.00263, p < 0.0001), confirming a role of cholinergic input in the more accurate intensity encoding of small cells at higher tone intensities (Fig 4). Chopper cells, which also receive cholinergic input (Fujino & Oertel, 2001), likewise showed reduced sound-evoked activity following atropine infusion, (75 dB SPL, α = 0.00263, p = 0.00098) but over a narrow intensity range. Atropine had no significant effect on the RLF of primary-like cells (75 dB SPL, α = 0.00263, p = 0.7419). These findings demonstrate that the primary action of cholinergic input on small cells and choppers is to increase sound-driven responses above 60 dB, resulting in a widened dynamic range that is essential for intensity coding.

**Fig 4:**
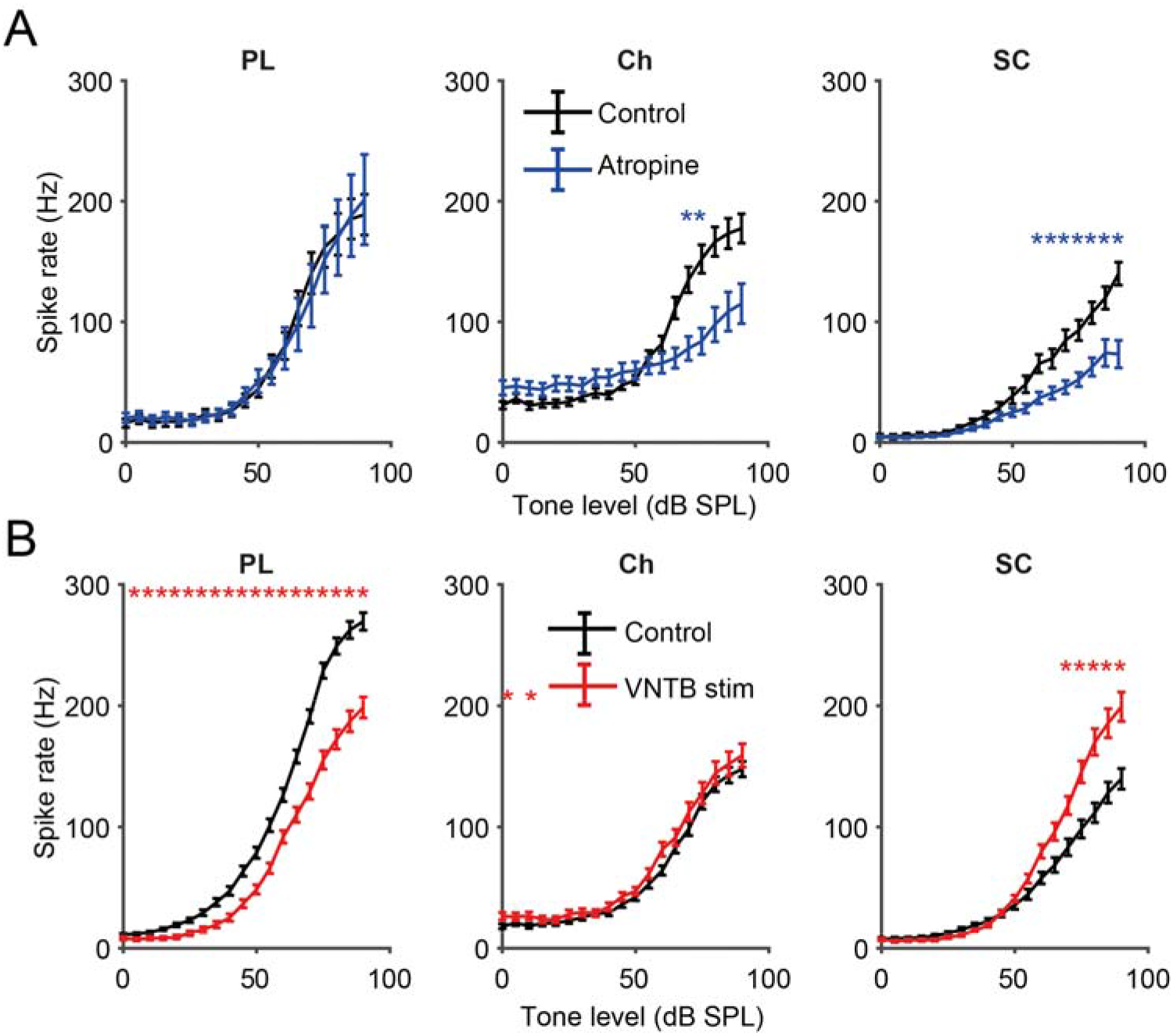
CN small cells receive excitatory cholinergic input from MOCs. A) Blocking muscarinic cholinergic receptors with atropine supresses CN small-cell and chopper responses to tones. Top row shows mean RLFs ± SEM for primary-like, chopper cells and small cells before and after local application of atropine via a drug-delivery probe. B) Electrically stimulating MOC neurons increases tone-evoked responses in small cells but not choppers. Mean RLFs ± SEM for primary-like, chopper cells and small cells are shown before and during MOC electrical stimulation. All statistical comparisons made using paired t-test with Bonferroni’s multiple comparison correction (α = 0.00263, stars signify p<0.05).

Next, to determine whether the MOCs were the key cholinergic modulators of small cells, we electrically stimulated MOC neuron somata in the VNTB while performing CN recordings. Successful targeting of MOC cells was confirmed by the reduction of CAP amplitudes during electrical stimulation. MOC stimulation significantly increased small cell firing rates in response to tones above 70 dB SPL but not to lower intensity or sub threshold tones (Fig 4, 75 dB SPL, α = 0.00263, p < 0.0001), suggesting that cholinergic MOC input to CN small cells enhances ANF-evoked activity. As expected, the overall effect of MOC activation on CN small cells was opposite to that achieved by cholinergic blocking. In contrast, VNTB stimulation had no effect on chopper firing (75 dB SPL, α = 0.00263, p = 0.461), while decreasing primary-like firing across the entire range of tone intensities (75 dB SPL, α = 0.00263, p < 0.0001). The reciprocal findings of increased firing rates with MOC stimulation and decreased firing rates with atropine in small cells confirm the role of MOC input to small cells in enhancing sound-driven responses. Remarkably, small cell excitation by MOC neurons can overcome the MOC suppression of cochlear output. The failure of MOC stimulation to alter firing rates in choppers suggests that the cholinergic effect observed originates rather from VNTB cells that are not MOCs.

### Small cells preserve higher-intensity tone coding in background noise

MOC projections to the cochlea suppress cochlear outer hair cell electromotility, thereby reducing cochlear amplification. This aids in unmasking of sounds from background noise, which facilitates the detection of tones and speech in noise (Dolan & Nuttall, 1988; Kawase & Liberman, 1993). Since small cells receive projections from MOC collaterals, we hypothesised that small cells are involved in encoding sounds in the presence of background noise. Signal-in-noise discrimination was tested in VCN neurons first by comparing RLFs to tones in silence with tones embedded in broadband noise.

As seen in individual and mean RLFs, small cells maintained their firing rates to medium to high intensity tones even in the presence of background noise (Fig 5, 75 dB p = 0.9677). In fact, small-cell spike rates in response to BF tones in the presence of 40- and 60 dB background noise were equivalent to their responses to tones in silence (Fig 6). Chopper units maintained their firing rates to BF tones in background noise at 40, but not 60 dB SPL (75 dB, p = 0.9677). In contrast, background noise degraded the responses to BF tones in primary-like cells, which showed increased rates and altered RLF patterns in the presence of 40- and 60 dB SPL broadband noise. These differences were not due to thresholds as they were apparent for cells of similar thresholds.

**Fig 5:**
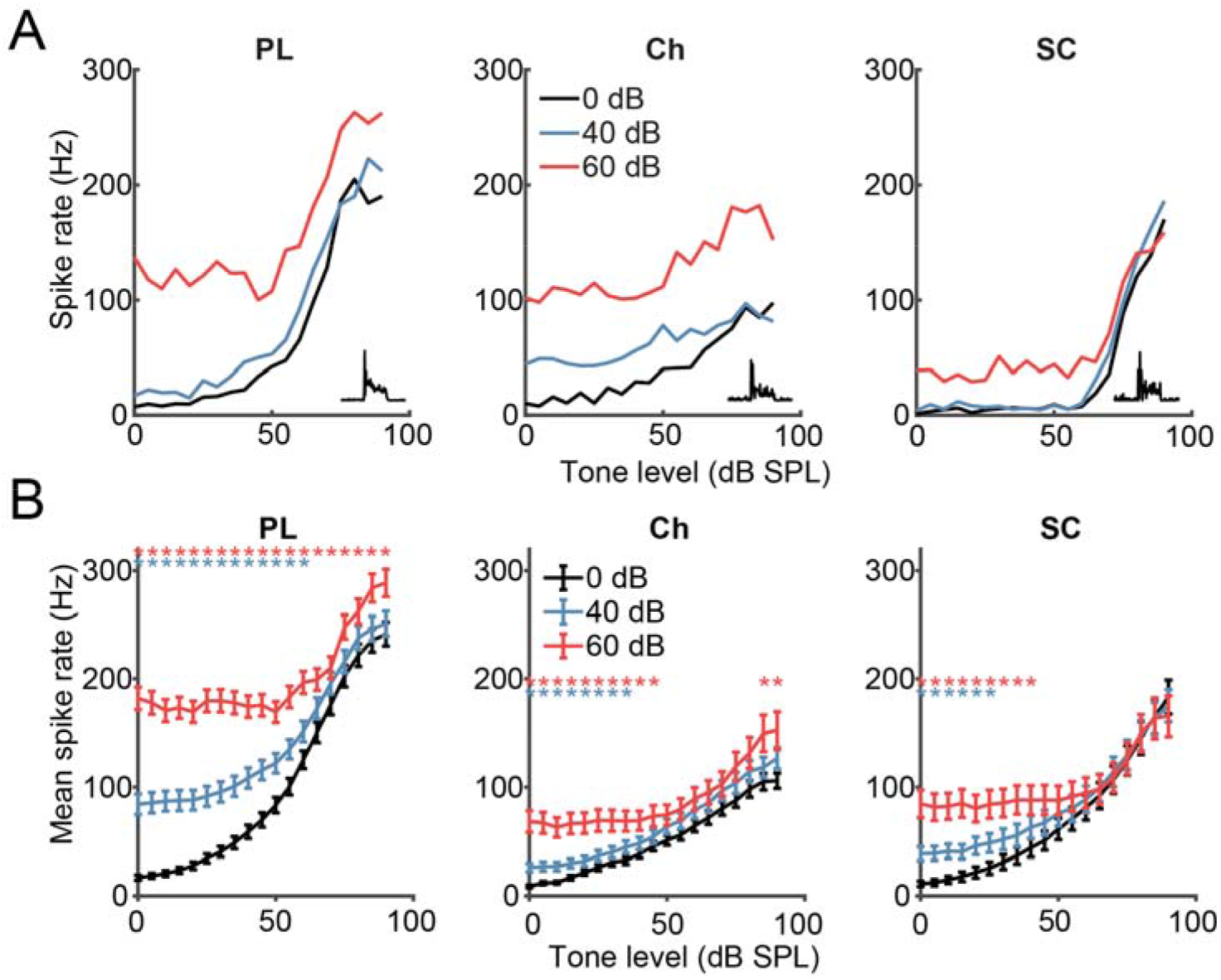
CN small cells maintain tone-level coding in the presence of background broadband noise. A) Single unit examples of RLFs in the presence of 0, 40- or 60 dB SPL background noise, for the 3 major cell-type classifications (columns). Insets show PSTH shapes. B) Mean (± SEM) RLFs of 3 major CN cell types in the presence of background noise.

**Fig 6:**
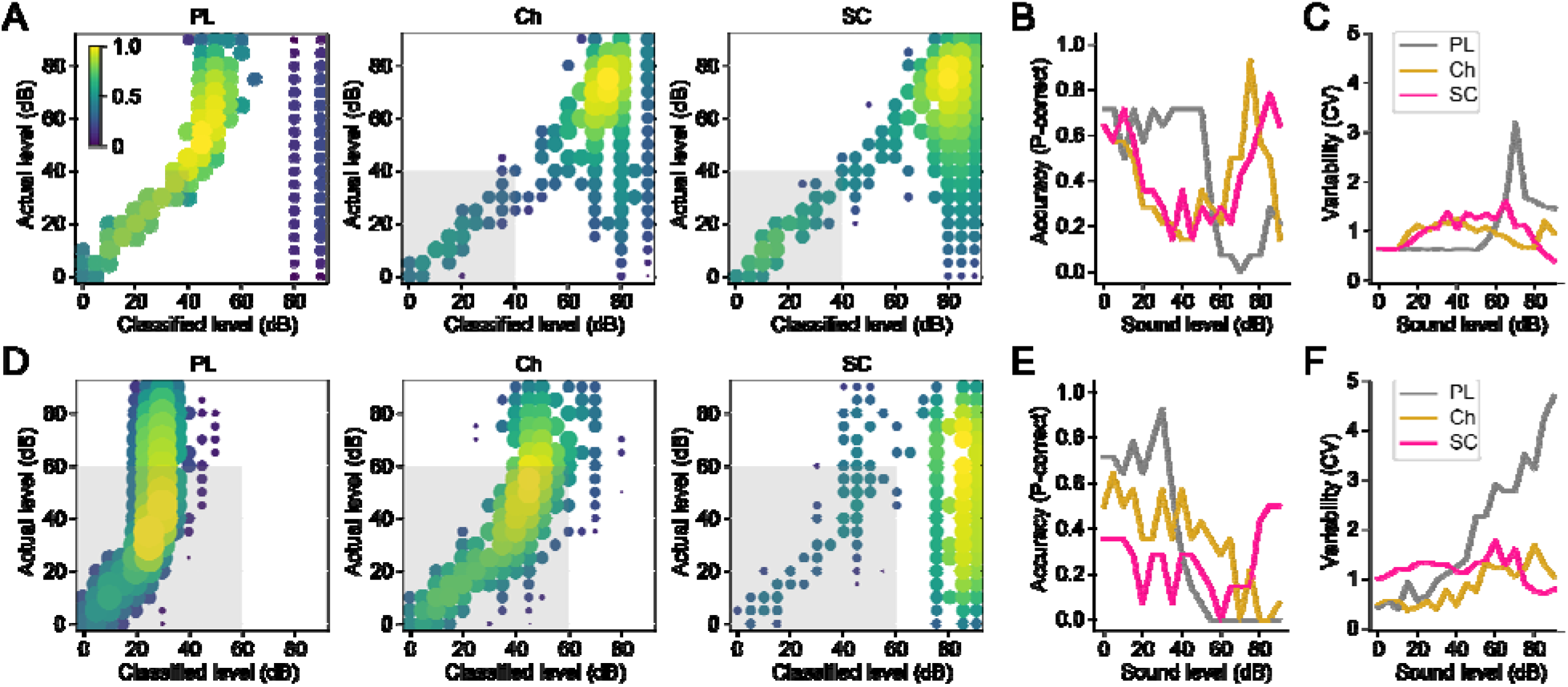
Small cells retain superior medium to high-level intensity coding in background noise. A) Representative PSTH-based classification of tone intensity for primary-like (PL), chopper (Ch), and small cells (SC) populations during constant 40 dB BBN. Colour bar = normalised kernel density across trials. B) Proportion of trials that show classified correctly (within ±5 dB of actual level) as a function of intensity compared among the three cell types. C) The variability of classification, or deviation from ‘correct’, measured by coefficient of variation (CV). D–F) Same format as (A–C) but for tone coding in 60 dB BBN. Gray boxes denote background noise levels.

To further examine intensity coding, we applied the PSTH-based classifier analysis on these same trials in both 40- and 60 dB broadband noise. Single-trial spike trains were classified based on templates generated in silence (Fig. 3D) in the same population of units. In primary-like cells during 40-dB background noise, we found that single-trial spike trains could not encode levels above 55 dB (Fig. 6A). Chopper and small cells retained coding above 65 dB with increased variability. The summary data showed that small cells are the most accurate coder with the lowest variability at the highest intensity tested (Fig. 6B–C; ANCOVA: F(1,2)=0.34, P=0.00085 across cell type). In 60-dB background noise, tone coding was nearly nonexistent for primary-like, and sparse above 60 dB for choppers (Fig. 6D). Only in small cells were high intensity coding above 75 dB preserved. At the highest intensity tested small cell retained 50% of accuracy with lowest variability among cell types (Fig. 6E–F; ANCOVA: F(1,2)=26.9, P=0.064). Taken together, these results demonstrate that, vis-à-vis VCN cell types, small cells reliably encode medium to high stimulus intensity in the presence of background noise.

### Small cells display precise envelope coding

Tracking sound envelopes is important for decoding human speech or animal vocalisations (Shannon et al., 1995). Here, we used amplitude modulated broadband noise (75 dB SPL) to evaluate envelope coding in CN neurons. Figure 7 (A-C) shows an example vector strength calculation for a single small cell at a modulation frequency of 32 Hz and depth of 12%, (Goldberg & Brown, 1969). This unit had a vector strength value of 0.43191, demonstrating accurate spike timing to the amplitude modulation. Figure 7 D shows that mean vector strength of the different VCN neurons peaked at a modulation frequency of 256 Hz, similar to that in previous studies (Sayles et al., 2013). At low modulation frequencies and depths, small cells had significantly greater vector-strength values than other VCN types (Fig 7) demonstrating superior envelope coding. For example, at 6% modulation depth and a modulation frequency of 16 Hz, Tukey’s multiple comparison test found that the mean value of small cells was significantly greater than choppers (p < 0.0001, 95% C.I. = 0.0437, 0.1345) and primary-like cells (p = 0.0014, 95% C.I. = 0.0172, 0.0877). The enhanced coding of low-frequency amplitude modulation in CN cells may be important for species-specific vocalisation processing, as guinea pig vocalisations contain predominantly low frequency amplitude modulation bursts (Berryman, 1976).

**Fig 7:**
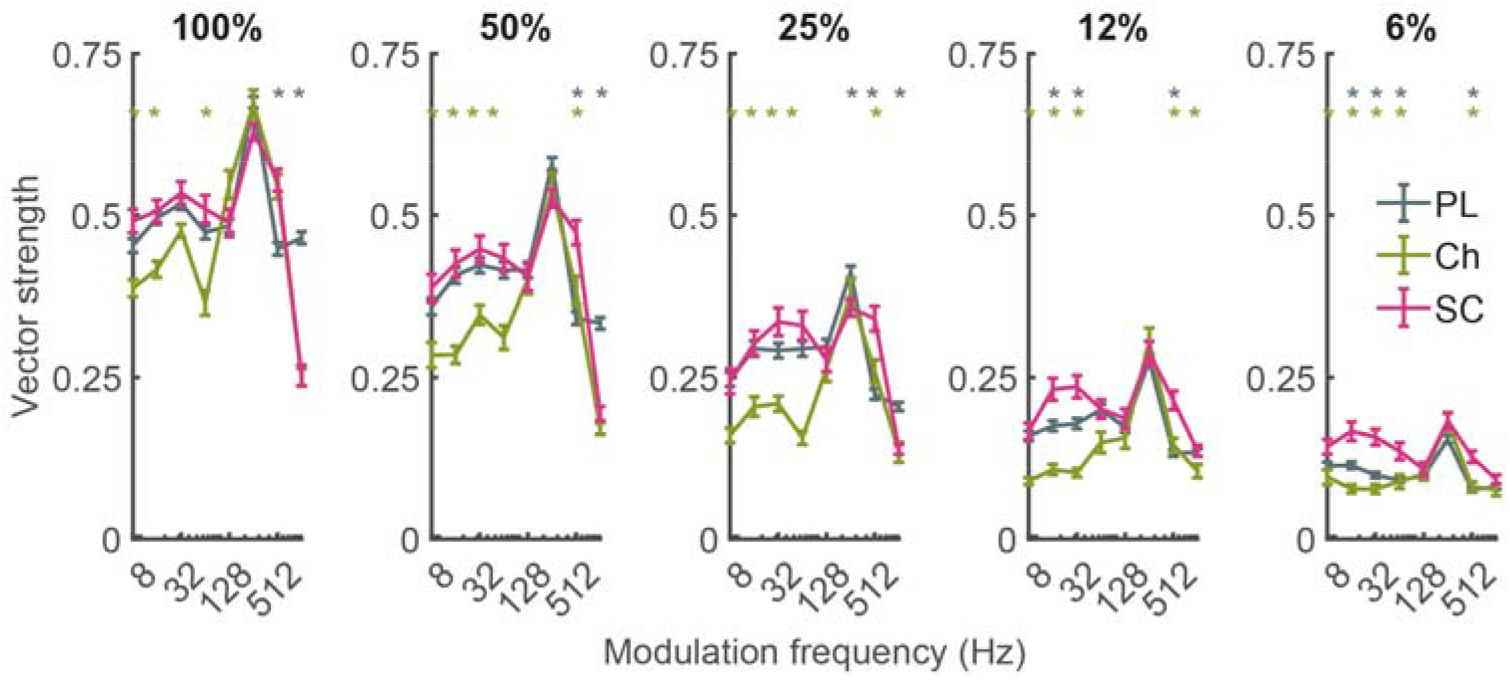
Small cells show more accurate amplitude-modulation coding than other VCN cell types. Modulation transfer functions show vector strength of spiking to sinusoidally modulated BBN at 5 different modulation depths (100-6%). CN small cells have significantly greater vector strength of firing at modulation frequencies below 128 Hz and at lower modulation depths. Stars indicate significance between CN small cells and other cell types (One-way ANOVA with Tukey’s multiple comparison test, p<0.05).

## DISCUSSION

Small cells receive exclusive afferent input from high-threshold, low/med SR ANFs, positioning them to encode sounds at levels well above threshold. Conversational speech (above 60 dB SPL) is encoded by temporal cues within amplitude modulated signals. Here, we showed that small cells encode both intensity and amplitude modulation with higher precision than other VCN cell types. For tones in background noise, small cells were the only VCN cell type that maintained coding above 60 dB SPL. The unique properties of small cells are likely mediated by cholinergic MOC input, which provide excitation to small cells capable of overcoming the suppressive effect of MOCs on the cochlea, thereby increasing their suprathreshold firing rates.

Small cells exhibited low SRs, high thresholds and steep, unsaturating (up to 90 dB SPL) rate-intensity slopes. These characteristics resemble the firing properties of the low/medium SR ANFs that innervate small cells and are similar to those identified in cells in the marginal region of the cat CN (Ghoshal & Kim, 1996, 1997). Those early studies also showed flat, bimodal or Unusual PSTH shapes, signifying a heterogenous population, similar to that shown here for guinea-pig CN small cells. However, in the present study, the difference between small-cell thresholds and other VCN cell types (∼5 dB) was less than the threshold difference between low/medium and high SR ANFs (∼20 dB; Liberman, 1978; Winter and Palmer, 1991), putatively due to the excitatory MOC cholinergic input to small cells demonstrated here. Blocking muscarinic, cholinergic receptors with atropine reduced small-cell spike rates while MOC stimulation, confirmed by CAP reductions, increased their firing rates. Both effects are consistent with the presence of excitatory cholinergic MOC input to the SCC. The greatest effect of blocking muscarinic cholinergic receptors was observed during suprathreshold sound stimulation, which would be expected to also activate the sound-driven, cholinergic projections from MOC collaterals to the SCC (Lee et al., 2006).

MOC collaterals to the CN can also influence other cell types (Baashar et al., 2019; Mulders et al., 2003, 2007) including T-stellate cells (planar multipolar cells with chopper responses) and D-stellate cells (radiate multipolar cells with onset chopper responses). Previous studies showed an increased firing rate in 94% of T-stellate cells following application of carbachol (AChR agonist) in vitro (Fujino & Oertel, 2001). However, Baashar et al. (2019) used tract tracing and double labelling to view contacts from MOC collaterals onto T- and D-stellate cells and demonstrated that MOC input to these CN cells was weaker than would be expected based on the early electrophysiological studies. Some studies have suggested that there is inhibitory MOC projection to the CN (Mulders et al., 2002), however the present study found no evidence for inhibitory MOC or cholinergic effect on CN small cells or choppers. One possible reason for this discrepancy is the more targeted stimulation of contralateral MOC neurons in the present study, compared to midline stimulation, (Mulders et al., 2002), which would activate all crossed olivocochlear fibers including the lateral olivocochlear system.

The present data reveal strong, excitatory MOC-collateral innervation of small cells in the SCC but weak input to T-stellate cells. Small-cell receptive fields in the present study had inhibitory sidebands, revealing a currently unknown source of inhibition. This could be the glycinergic, D-stellate cells, which provide wideband inhibition to several cell types in the CN (Arnott et al., 2004; Oertel et al., 1990). However, further studies are required to confirm this pathway. An alternative source of inhibition is the L-stellate cell, which also responds to carbachol and thus potentially of olivocochlear origin (Ngodup et al., 2020). However, prediction of sources based on bath infusion of cholinergic modulators must be made with caution, due to multiple sources of cholinergic modulation to the CN (Mellott et al., 2011).

We demonstrated that small cells more accurately encode tones-in-noise compared to other VCN cell types, putatively due to their restricted ANF inputs and unique input from MOC collaterals. Activating the MOC pathway to the cochlea suppresses the cochlear amplifier, benefitting speech-in-noise processing (de Boer et al., 2012; Winslow & Sachs, 1987). The function of MOC collaterals to the SCC may be to further aid speech coding. Small cells have the characteristics necessary for superior suprathreshold sound encoding and accurately encode stimulus intensity (Figs 3&6). Pre-emptive MOC-pathway enhancement protects the cochlea from synaptopathic acoustic trauma (Boero et al., 2018), but whether increased MOC activation following synaptopathic noise exposure could reverse speech-in-noise deficits is still unknown. Conversely, in humans with tinnitus or hyperacusis, an increased contralateral suppression of otoacoustic emissions (Knudson et al., 2014) suggests an over-responsive MOC system in that pathology. MOC neurons are modulated by descending input from the inferior colliculus (Malmierca et al., 1996; Robertson & Mulders, 2000; Schofield & Cant, 1999), auditory cortex (Coomes & Schofield, 2004; Mulders & Robertson, 2000), and local inhibitory interneurons of the medial nucleus of the trapezoid body (Torres Cadenas et al., 2020). Descending modulation may be a route for auditory attention to affect peripheral processing, which may explain attentional facilitation of speech-in-noise understanding (de Boer et al., 2012). Descending modulation of MOC activity would also affect MOC collateral input to CN small cells, which accurately encode intensity, providing an additional action pathway for auditory attention.

Hearing damage in the absence of threshold shifts preferentially affects the higher threshold ANFs (Furman et al., 2013) that provide the sole afferent input to small cells. Cochlear synaptopathy is a potential major health issue in humans, as human temporal bones demonstrate widespread synaptic damage (Viana et al., 2015). One downstream effect of cochlear synaptopathy may be impaired speech perception, especially in the presence of background noise. This widespread complaint has been attributed to cochlear synaptopathy, but this relationship is still unclear (Plack et al., 2014). Human studies suggest that such perceptual deficits reflect difficulties in coding temporal envelopes (Bharadwaj et al., 2014), expected to impair speech recognition (Shannon et al., 1995). Temporal-coding deficits after synaptopathy are not apparent in ANFs (Heeringa et al., 2020), suggesting their origin may be central. The enhanced intensity coding and precise envelope coding shown here for small cells primes the SCC as an important subcortical area for speech processing is expected to be preferentially targeted by cochlear synaptopathy. In humans, the SCC occupies a large proportion of the CN and is therefore poised to play a major role in central mechanisms of auditory neuropathy (Moore & Osen, 1979). Speech coding in human small cells may be impaired following cochlear synaptopathy, and modulation of CN activity via the MOC pathway may be a potential treatment for the associated speech-in-noise deficits.

## Acknowledgements

The authors would like to thank Mr. Dileepkumar for engineering support throughout this project. This study was supported by National Institutes of Health grant R01-DC017119 (S.E.S.).

---

**Fig S1:**
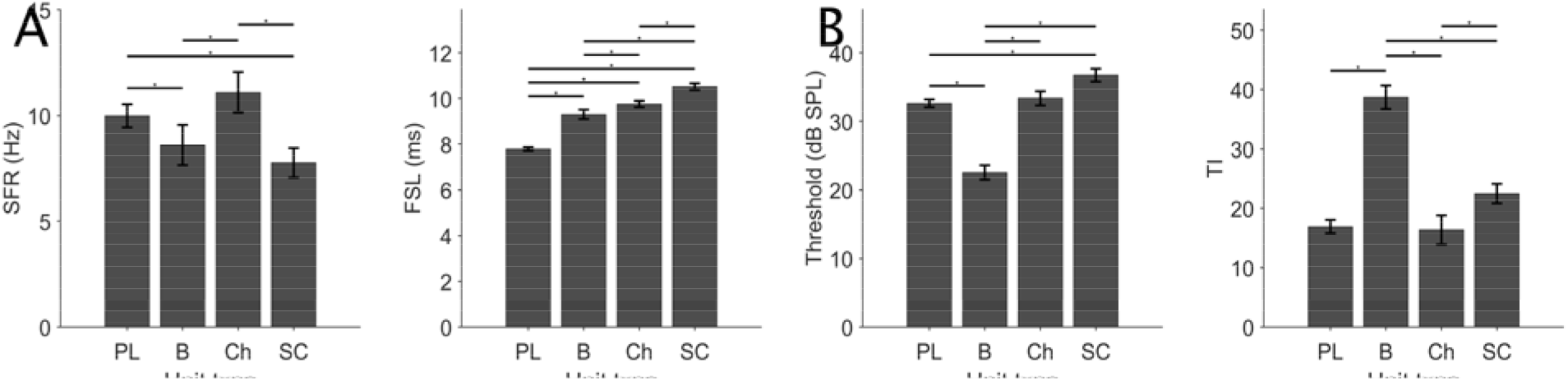
Small cells show lower SFRs and longer FSLs than other VCN units. A) Bar plot (mean ± SEM) of SFR for CN cell types (primary-like (PL), chopper (Ch), small cell (SC) and buildup. B) Bar plot (mean ± SEM) for FSL. C) Bar plot (mean ± SEM) of thresholds for each CN cell type. D) Bar plot (mean ± SEM) of total inhibitory (TI) area of the receptive field. Comparisons between groups were made using a Kruskal-Wallis test with Tukey’s multiple comparison test *p<0.05.

**Table S1:**
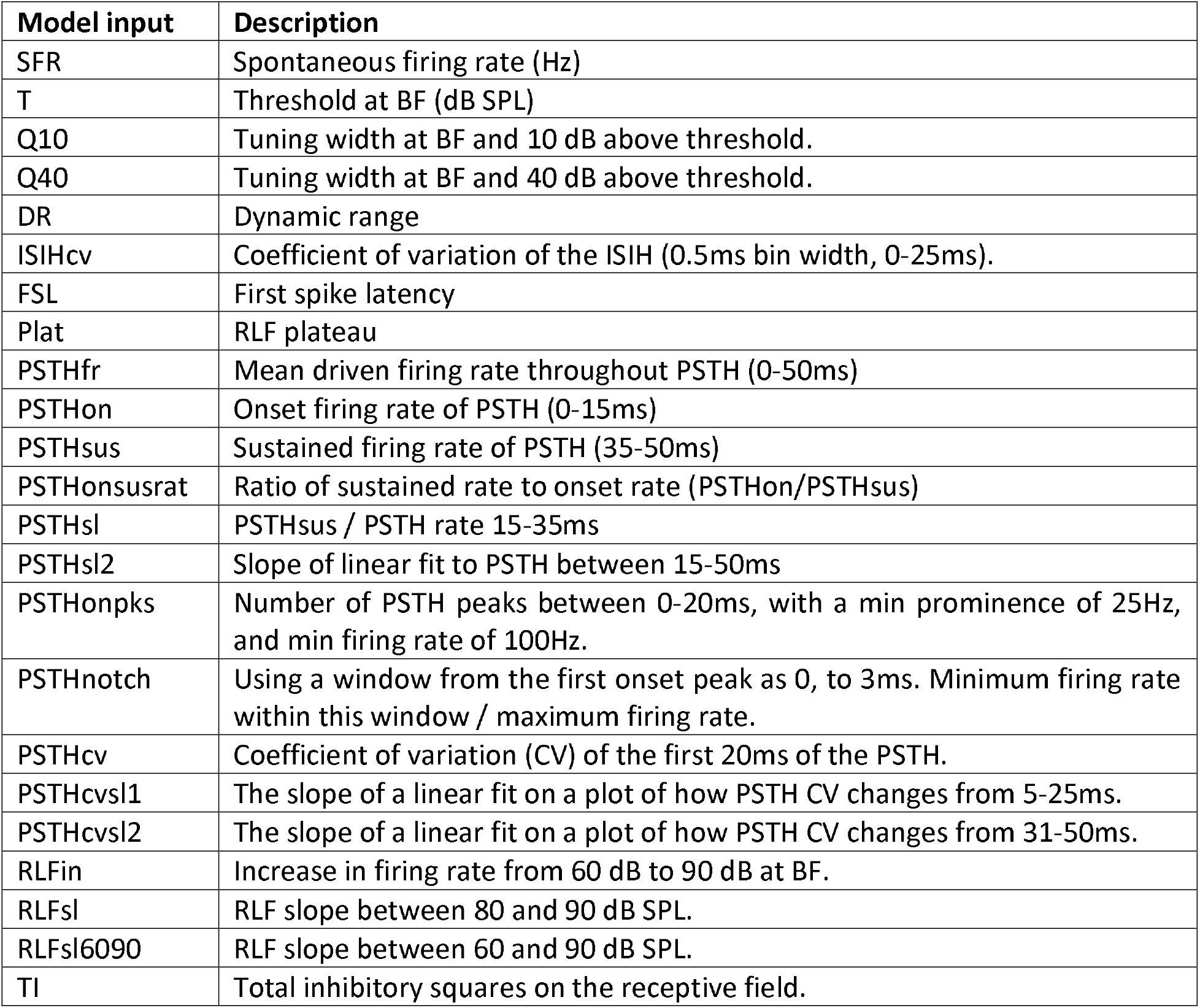
Inputs for machine learning CN cell typing model.

